# Quantification of human-mosquito contact rate using surveys and its application in determining dengue viral transmission risk

**DOI:** 10.1101/821819

**Authors:** Panpim Thongsripong, Zhuolin Qu, Joshua O Yukich, James M Hyman, Dawn M Wesson

**Affiliations:** Department of Tropical Medicine, Tulane University, New Orleans, Louisiana, United States of America; Department of Mathematics, Tulane University, New Orleans, Louisiana, United States of America

**Author notes:** Microbiology Department, California Academy of Sciences, San Francisco, California, United States of America (Panpim Thongsripong).

## Abstract

*Aedes*-borne viral diseases, including dengue fever, chikungunya, and Zika, have been surging in incidence and spreading to new areas where their mosquito vectors thrive. To estimate viral transmission risks, availability of accurate local transmission parameters is essential. One of the most important parameters to determine infection risk is the human-mosquito contact rate. However, this rate has rarely been characterized due to the lack of a feasible research method. In this study, human-mosquito contact rates were evaluated in two study sites within the Greater New Orleans Region by asking a group of survey participants to estimate mosquito bites they experienced in the past 24 hours. The fraction of the mosquito bites attributed to *Ae. aegypti* or *Ae. albopictus* was estimated by human landing sampling. The results showed a significantly higher outdoor mosquito bite exposure than indoor exposure. The number of reported mosquito bites was positively correlated with the time that study participants spent outside during at-risk periods. There was also a significant effect of the study site on outdoor bite exposure, possibly because of the difference in the numbers of host-seeking mosquitoes. We use a mathematical dengue virus transmission model to estimate the transmission risks in the study areas based on local conditions. This compartmental model demonstrated how the observed difference in the human-*Aedes* contact rates in the two study sites would result in differential dengue transmission risks. This study highlights the practicality of using a survey to estimate human-mosquito contact rates and serves as a basis for future evaluations. Combined with the use of mathematical modeling, this innovative method may lead to more effective mosquito-borne pathogen prevention and control.

**Author summary:** Even though the human-mosquito contact rate is among the most important indicators of mosquito-borne viral transmission risk, it is rarely characterized in the field. Human Landing Capture is a gold standard method to quantify this rate, but it ignores variables such as human behaviors and lifestyles. In this study, we tested the feasibility of using surveys to quantify mosquito bite exposure in the Southern United States. The survey results, combined with mosquito species proportion data, were used to estimate the contact rate. These rates are key parameters used in mathematical models to determine transmission risks. We found that bite exposure occurred more often outside homes and people who spent more time outdoors in the evening and night had a higher exposure. Our model analysis shows that the human-mosquito contact rate is one of the most important parameters determining outbreak potential. Disease control programs should focus their efforts on reducing this rate in addition to the mosquito density. Future studies should test if the entomological contact rates described by surveys correlate with disease incidences or other entomological indices. This study highlights the importance of characterizing how vector-human contact rates may respond to changing human behaviors and environments.

## Introduction

Globally, mosquito-borne viral diseases are on the rise. In the past few decades, diseases such as dengue, West Nile fever, chikungunya, and Zika have emerged and persisted in the parts of the world where their mosquito vectors thrive [1–5]. It has been estimated that hundreds of thousands of people die from mosquito-borne diseases each year [6]. Population growth, unplanned urbanization, global warming, intercontinental travel, and the breakdown of mosquito control infrastructure have all contributed to the expansion of mosquito vectors in multiple locations throughout the world [7–11].

Dengue fever is the most common and widespread mosquito-borne viral disease in the world [12]. According to a recent study [13] about 390 million dengue viral infections occurred in 2010, higher than the 50-100 million previously estimated by the World Health Organization. Four serotypes of dengue virus (DENV) can be transmitted by two species of mosquito vectors in the genus *Aedes*: *Aedes aegypti* (Linnaeus), the Yellow Fever mosquito, and *Aedes albopictus* (Skuse),the Asian Tiger mosquito [14]. Both species are highly anthropophilic [1]. They are widespread in residential settings of tropical and sub-tropical parts of Asia, Latin America, Africa, and the Pacific [15]. Because of a suitably warming climate and the availability of larval habitats, both *Aedes* species have gained a foothold in the Southern United States and Southern Europe [16–18]. This, in combination with increases in international travel, results in a possibility that DENV may emerge in these areas.

Mathematical models can help guide the design of effective preemptive and ongoing disease control programs [19]. For these models to be effective, they require accurate estimates of local transmission parameter values. One of the most important parameters determining pathogen transmission is the human-mosquito contact rate, which we define as the total number of times humans are bitten by mosquitoes each day in the area of interest [20]. Unfortunately, these rates are rarely characterized because of the lack of appropriate research methods. The paucity of contact rate data hinders our progress in understanding how changing environments and human behaviors will affect mosquito-borne virus transmission and emergence. We need to know how often, and under what circumstances, humans are exposed to mosquito bites to plan effective mitigation strategies.

To date, only a few approaches have been used to approximate contact patterns in the field. Human Landing Capture (HLC) is the traditional gold standard method to monitor human-vector contact patterns in malaria transmission [21, 22]. This method involves human volunteers collecting mosquitoes that land on them to feed, typically at night when *Anopheles* spp., the vectors transmitting malaria, seek a blood meal. A well-designed HLC study could potentially approximate the contact rate when humans are bitten by mosquitoes while sleeping. However, because *Aedes* spp. bite during the day when humans could actively interrupt or avoid mosquito bites, this could result in a potential bias for the HLC estimates. The contact rate depends heavily on housing infrastructure, human behaviors, and lifestyle differences that cannot be captured easily by an HLC experiment [23–25].

In this study, we approximated the contact rates between *Aedes* spp. and humans in the Greater New Orleans Region using a questionnaire-based survey and a small-scale HLC experiment. A short questionnaire in the form of door hangers was used to ask research participants about the frequency and location of mosquito bite exposures in the past 24 hours. An HLC study was performed to determine the proportion of mosquito bites that belong to either *Ae. aegypti* or *Ae. albopictus*. Next, the contact rates between humans and the *Aedes* species were calculated. Finally, a deterministic compartmental SEIR (Susceptible, Exposed, Infected, Recovered) model describing DENV transmission by *Ae. aegypti* and *Ae. albopictus* was used to compare how the model predictions depend on the locally characterized human-mosquito contact rates from two distinct locations.

The ultimate goals of this study were: 1) to test the feasibility of using questionnaire-based surveys to quantify human-mosquito contact rates, 2) to understand how environmental factors and human behaviors may impact mosquito bite exposure, and 3) to model how changes in human-mosquito contact rates impact pathogen transmission outcomes.

## Methods

### Study sites and survey methods

We designed two questionnaires in the form of door hangers. We intentionally designed short questionnaires to encourage participation. The first questionnaire (Supplementary Data 1) was used in a preliminary survey to explore the range of bite exposure and to estimate the return rate. The research participants were asked to indicate the number of mosquito bites they received within the past 7 days, the locations in which they experienced mosquito bites most often, and the frequency of mosquito bite exposures inside homes. All questions in this questionnaire were in a multiple-choice format.

The second questionnaire (Supplementary Data 2) was designed after the preliminary survey. The questions included open-ended questions inquiring about the amount of mosquito bites participants received both indoors and outdoors, where they had received the outdoor bites, and the time spent outside in the past 24 hours. The questionnaire also collected demographic data including age range, gender, and number of people in their household.

In the preliminary survey, the questionnaires were distributed in August and September of 2016 in three study sites: the Bywater and 7^th^ Ward neighborhoods of Orleans Parish (ORL), the Bridge City neighborhood of Jefferson Parish (JEF), and the Oak Harbor and Eden Isle neighborhoods of St. Tammany Parish (TAM). Four street blocks were randomly chosen per month from each of the three study sites. The questionnaires were distributed to all addresses in the chosen blocks and collected back the next day.

In the second survey, only two study sites, ORL and TAM, were included. The study period was from April to August 2017. In each month, 4 street blocks from each study site were randomly selected, without replacement, to receive the questionnaires on Sundays, and another 4 blocks on either Wednesdays or Thursdays. The questionnaires were distributed to all addresses in the chosen blocks and retrieved back the next day. No identifying information or addresses were collected from the study subjects, and the Tulane University’s Internal Review Board (IRB) approved the full-review exempt status of both surveys (IRB reference number: 16-923467E).

The ORL site was in an urban environment close to New Orleans city’s downtown area. Compared to the other two study sites, ORL’s residents were younger and lived in a smaller household (almost 40% of all households were a 1-person household; US 2010 Census). Its population median age was 38 (40 in JEF, and 50 in TAM; US 2010 Census). ORL was a racially mixed neighborhood (52.85% African American and 41.86% White; US 2010 Census). JEF and TAM are located further away from the city’s downtown area in a more sub-urban environment. TAM had the highest average household income ($96,415; 2016 ACS 5-year estimates) compared to ORL ($55,709), and JEF ($49,928). TAM also had the highest percentage of households that were classified as “Family Household” (76.40%; US 2010 census). Racial diversity was lowest in TAM (89.18% of total population were White). The population variables of the study sites are shown in detail in Supplementary Table 1.

### Human Landing Catch (HLC)

HLC experiments were performed in ORL and TAM to investigate the species composition of host seeking mosquitoes from April to August 2017. Two locations were chosen from each study site. In each location and month, HLC was performed once in the morning and once in the evening on two separate days. Each collection consisted of two 45-minute capturing sessions with an up to 15 minute break in between. The morning collection started within 30 minutes after sunrise, and the evening collection stopped within 30 minutes before sunset. The HLC locations were shaded outdoor areas. The collector was seated on a chair with the legs exposed from the shoes up to the knees, and the lower arms were exposed from the elbows down. Collection of landing mosquitoes from the collector’s own body was done using a portable aspirator and the mosquitoes were either identified on site, when possible, or transported back to the laboratory for further identification using a microscope. A single collector took part in all the HLC sessions.

### Survey and HLC data analysis and statistical tests

Because the first survey was a preliminary data collection with a small sample size, only the data from the second survey was analyzed with statistical tests. In the second survey, the sampling method was a two-stage stratified cluster sampling. To account for the differential probabilities of selection due to the study design and to ensure more accurate estimates, a sampling weight for each participant was calculated based on the selection probability proportional to size. The population cohort was defined as persons aged >18 years old who lived in the two study sites at the time of sampling. The Primary Sampling Unit (PSU) was at a residential block level. The sampling probability of each block was 1/B_i_, where B_i_ is the total number of blocks in study site i. The Secondary Sampling Unit (SSU) was at the research participant level. The probability that a person in each household was selected was 1/P_j_, where P_j_ was the household size for address j.

All data analysis was done using R (version 3.3.3) and R studio. The data and weights were defined to create a Survey Object using Survey package [26]. Sampling weight for each data point was calculated as the inverse of the probability of selection. Specifically, weight for each data point was equal to (1/B_i_+1/P_j_)^-1^. All statistical tests downstream of the weighting procedure were analyzed with the functions within Survey package. To compare the numbers of reported bites and the time spent outside within the past 24 hours between groups, Wilcoxon Rank Sum tests were used. Spearman’s correlation tests were used to determine the correlation between the time spent outside at each time interval and the numbers of reported bites received outdoors.

Two generalized linear models assuming quasi-Poisson distribution as the probability distribution function of the response variable, with log link function, were created to analyze the data. The first model used the total time spent outside between 5 pm to 6 am (evening and nighttime) as a response variable. In this model, the independent variables included the age range and gender of research participants, weekend/weekday setting, and study sites. The second model used numbers of reported bites received outdoors within the past 24 hours as a response variable. The independent variables included in this model were the time spent outside within the past 24 hours, the gender of research participants, the month of data collection, and the weekend/weekday setting.

For HLC data analysis, comparisons between the numbers of landed *Ae. aegypti* or *Ae. albopictus* between study sites and between times of collection were determined using the Wilcoxon Rank Sum test. The proportions of *Ae. aegypti* and *Ae. albopictus* from HLC were calculated based on average values of landing mosquito types across all HLC sessions for both study sites.

### Dengue epidemiological compartmental model description and assumption

Our compartmental mathematical model described the transmission of one serotype of DENV by both vector species: *Ae. aegypti* and *Ae. albopictus*. We used this model to estimate and predict quantities of interest at the initial epidemic spread. This model was adapted from a mathematical mosquito-borne disease model published in a study by Manore *et al.* [20]. The human-mosquito contact rates used in the model were based on the local survey data. We defined human-mosquito contact rate (B) as the number of biting events that occurred by all mosquitoes of a given species on the human population in the area of interest within a 24-hour period. In other words, it was the number of bites all humans in the area of interest received from that mosquito species within 24 hours. Note that we defined the mosquito’s *biting rate* as a *per capita* rate of bites that a typical single mosquito may give to humans per unit time. As a result, a mosquito’s biting rate was different from a human-mosquito contact rate.

The human population was divided into 4 compartments: susceptible (S_h_), exposed (E_h_), infectious (I_h_), and recovered/immune (R_h_). The *Ae. aegypti* mosquito population was divided into 3 compartments: susceptible (S_g_), exposed (E_g_), and infectious (I_g_). The *Ae. albopictus* mosquito population was also divided into 3 compartments: susceptible (S_b_), exposed (E_b_), and infectious (I_b_). The total population sizes for *Ae. aegypti*, *Ae. albopictus* and humans were N_g_ = S_g_ + E_g_ + I_g_, N_b_ = S_b_ + E_b_ + I_b,_ and N_h_ = S_h_ + E_h_ + I_h_ + R_h_, respectively. We assumed that the two vector species do not interact. This means, for example, that the carrying capacities of the two species were independent from each other. Supplementary Figure 1 shows a diagram of the model including the relationship among all population compartments.

Humans entered the susceptible class S_h_ with a per capita birth rate Ψ_h_. Humans were bitten by *Ae. aegypti* with a rate of B_g_/N_h_ (bites per person per day) or by *Ae. albopictus* with a rate of B_b_/N_h_. These biting *Ae. aegypti* or *Ae. albopictus* had a probability of I_g_/N_g,_ or I_b_/N_b,_, of being infectious, respectively. If a mosquito was infectious, then there was a probability of β_h_ that the person will become infected. When a human was infected, they moved from susceptible class S_h_ to the exposed class E_h_. After an average intrinsic incubation period of 1/ν_h_ days, they moved to the infectious class I_h_. Humans in the infectious class can infect other mosquitoes upon contacts. After an average recovery time 1/γ_h_ days, the infectious humans recovered and moved to class R_h_. Recovered persons were assumed to have immunity to the infecting DENV serotype for the entire period of the simulation. In addition, humans of all status left the population through a per capita natural death rate μ_h_. The death rate due to disease was assumed to be very low and negligible. The human population size was assumed to be stable (Ψ_h_ = μ_h_), and migration of mosquitoes and humans was low and negligible.

When a susceptible *Ae. aegypti* mosquito bit humans at a biting rate of B_g_/N_g_ (bites per mosquito per day), there was a probability I_h_/N_h_ that the persons being bitten were infectious. If the person was infectious, then the biting *Ae. aegypti* mosquito in the class S_g_ became infected with a probability β_g_ and moved to the exposed class E_g_. After an average extrinsic incubation period 1/ν_g_ days, the mosquito advanced to the infectious class I_g_. Similarly, when a susceptible *Ae. albopictus* mosquito bit humans at a biting rate of B_b_/N_b_, there is a probability I_h_/N_h_ that the persons were infectious and a probability β_b_ that the mosquito became infected and advanced to the exposed class E_b_. After an extrinsic incubation period 1/ν_b_ days, the *Ae. albopictus* mosquito advanced to the infectious class I_b_. Both mosquito species remained infectious for life.

Female mosquitoes entered the susceptible class through recruitment from the pupal stage. The recruitment term for mosquitoes was proportional to the egg-laying rate of adult female mosquitoes and accounted for the hatching rate of eggs and survival of larvae and pupae. The aquatic stages were not explicitly included in the model and were approximated by a density-dependent recruitment (birth) rate. We assumed that all adult female *Ae. aegypti* and *Ae. albopictus* mosquitoes had the same per capita natural death rate μ_g_ and μ_b_, respectively. In this model, dengue infection did not affect the mosquito death rate or biting rate.

### Model equations

Our ordinary differential compartmental equations modeling dengue transmission were:

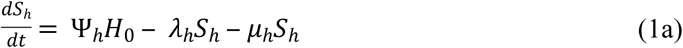

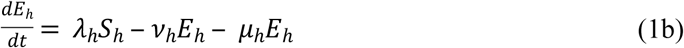

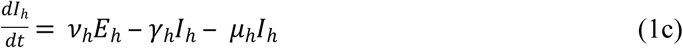

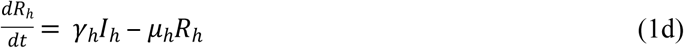

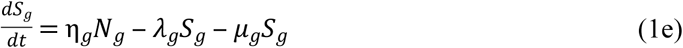

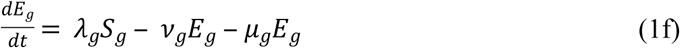

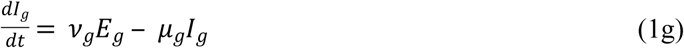

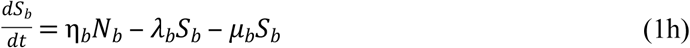

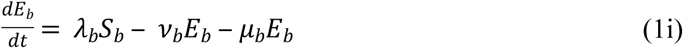

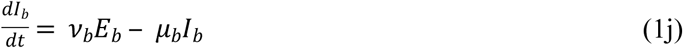

The female *Ae. aegypti* and *Ae. albopictus* recruitment rates were:

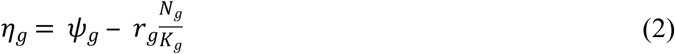

and

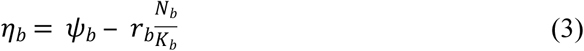

Here, Ψ_g_ and Ψ_b_ were the per capita natural birth rates of female *Ae. aegypti* and *Ae. albopictus*, respectively. In the absence of density dependence, r_g_ and r_b_ were the intrinsic growth rates of female *Ae. aegypti* and *Ae. albopictus*, respectively, where r_g_ = Ψ_g_ – μ_g_ and r_b_ = Ψ_b_ – μ_b_. K_g_ and K_b_ were the carrying capacity of the female *Ae. aegypti* and *Ae. albopictus*, respectively, in the area of interest.

The force of infection from mosquitoes to humans (λ_h_) was the product of the average number of bites a person received from mosquitoes per day (B_g_/N_h_ and B_b_/N_h_), the probability that the mosquito was infectious (I_g_/N_g_ and I_b_/N_b_), and the probability of virus transmission from the biting and infectious mosquito to the human (β_h_),

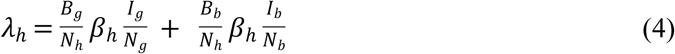

The force of infection from humans to *Ae. aegypti* and to *Ae. albopictus* (λ_g_ and λ_b,_ respectively) were the product of the number of bites per mosquito per day (B_g_/N_g_ and B_b_/N_b_, respectively), the probability that the bitten human was infectious (I_h_/N_h_), and the probability of pathogen transmission from an infected human to the biting mosquito (β_g_ and β_b_, respectively).

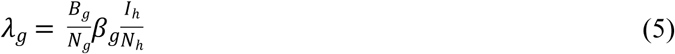

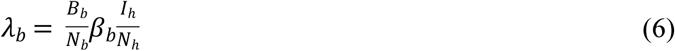

### Model parameters

The contact rates of humans and *Ae. aegypti* (B_g_) or *Ae. albopictus* (B_b_) were obtained from this study. Other parameters were obtained from other sources (Table 1).

**Table 1.** Model parameters, their baseline values and ranges, and sources [20,27–34].

|  | Parameter | Unit | Value | Range | Source |
| --- | --- | --- | --- | --- | --- |
| $H_0$ | Human population size, ORL | Human | 10,157 | - | US census 2016 estimates |
|  | Human population size, TAM |  | 7,385 | - |  |
| $B_g$ | *Ae. aegypti*-human contact rate, ORL | Day<sup>-1</sup> | 26,389 | 17,094 - 35,684 | from this study |
|  | *Ae. aegypti*-human contact rate, TAM |  | 5,484 | 3,777 - 7,197 |  |
| $B_b$ | *Ae. albopictus*-human contact rate, ORL | | 40,916 | 26,504 - 55,329 | |
|  | *Ae. albopictus*-human contact rate, TAM |  | 9,834 | 6,773 - 12,895 |  |
| $\beta_h$ | Probability of transmission from mosquito to human given an infectious bite | - | 0.33 | 0.10 - 0.75 | [20,27] |
| $\beta_g$ | Vector competence for *Ae. aegypti* | - | 0.25 | 0.03 - 0.76 | [28,29] |
| $\beta_b$ | Vector competence for *Ae. albopictus* | - | 0.06 | 0.01 - 0.56 | [28,29] |
| $1/v_g$ | EIP for *Ae. aegypti* | Day | 6.5 | 2 - 33 | [30] |
| $1/v_b$ | EIP for *Ae. albopictus* | | | | |
| $1/v_h$ | IIP | Day | 6 | 3 - 10 | [31] |
| $\Psi_g$ | Per capita recruitment rate of *Ae. aegypti* | Day<sup>-1</sup> | 4.93 | 3.89 - 5.97 | [32] |
| $\Psi_b$ | Per capita recruitment rate of *Ae. albopictus* | | | | |
| $K_g$ | Carrying capacity of *Ae. aegypti* | Mosquito | $10H_0$ | $3H_0 - 17H_0$ | Estimated |
| $K_b$ | Carrying capacity of *Ae. albopictus* | | | | |
| $1/\gamma_h$ | Viremic period in human | Day | 5 | 4 - 14 | [31,33] |
| $\mu_h = \Psi_h$ | Per capita death and birth rate for human | Year<sup>-1</sup> | 1/75.7 | 1/74.9 - 1/81.3 | CDC's wonder database |
| $\mu_g$ | Per capita death rate for *Ae. aegypti* | Day<sup>-1</sup> | 1/18 | 1/11 - 1/55 | [32,34] |
| $\mu_b$ | Per capita death rate for *Ae. albopictus* | | | | |

The bite number, ρ_h_, was the total number of bites a typical human received per person per day, regardless of mosquito species, and was estimated from our survey. The proportion of bites, p_v_, that belonged to mosquito species v was estimated from HLC data. The number of mosquito bites that belonged to mosquito species v that humans received per person per day (or the bite exposure rate) was

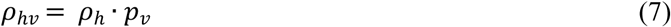

If H_0_ was the human population size, then, the number of mosquito bites from mosquito species v that all humans in the population received per day (or the contact rate) is

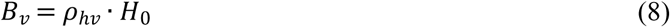

### The basic reproductive number (R_0_)

The calculations and model analyses were done in MATLAB R2018a (version 9.4.0). The model outcomes of interest were 1) the initial rate of disease spread by evaluating the basic reproduction number (R_0_) and 2) the initial transient disease dynamics by evaluating the timing and magnitude of the first epidemic peak. Disease-free equilibrium points are steady-state solutions where there is no disease; i.e., no exposed or infectious individuals for both humans and mosquitoes. Let *X = (N_h_, E_h_, I_h_, R_h_, N_g_, E_g_, I_g_, N_b_, E_b_, I_b_),* then the model for dengue transmission had exactly one disease-free equilibrium point, *X_dfe_ = (H_0_, 0, 0, 0, K_g_, 0, 0, K_b_, 0, 0)*, with no disease in the population.

In a homogeneously mixed population, the basic reproduction number (R_0_) is the expected number of secondary infections that one infectious individual would cause over the duration of the infectious period in a fully susceptible population [35]. From this definition, it can be logically interpreted that when R_0_ < 1, each infectious individual produces less than one new infected individual on average and the pathogen transmission ‘dies out’ from the population. Conversely, if R_0_ > 1, the pathogen is able to invade the susceptible population.

The next generation operator approach was used to calculate R_0_ [36]. The description of the calculation of R_0_ using the next generation operator is described in detail in Appendix A, which resulted in R_0_ expression:

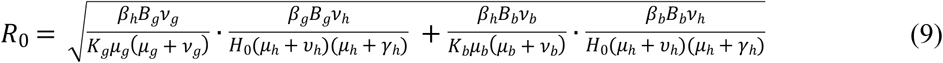

In a fully susceptible human population, the number of new human infections caused by one infected *Ae. aegypti*, or the basic reproductive number for the disease transmission from *Ae. aegypti* to human, was

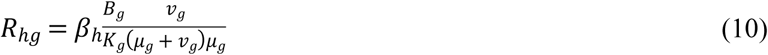

In this expression, 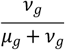 was the probability of *Ae. aegypti* surviving the exposed stage and becoming infectious. 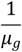 was the lifespan of *Ae. aegypti.* The product of these two terms, or 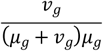, equaled to the average number of days that *Ae. aegypti* was infectious. As a result, R_hg_ can be seen as the product of 1) the number of bites per day per mosquito, or 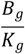, 2) the probability of a successful transmission per bite, or *β_h_*, and 3) the number of days in the infectious period, or 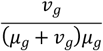.

Similarly, the basic reproductive number for the disease transmission from *Ae. Albopictus* to human, from human to *Ae. aegypti*, and from human to *Ae. albopictus*, respectively, was

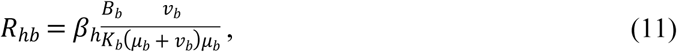

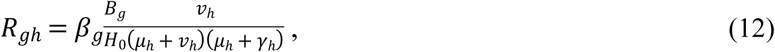

and

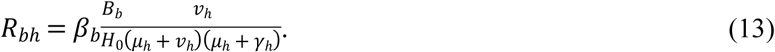

The basic reproduction number R_0_ in (9) can be expressed in terms of these quantities as

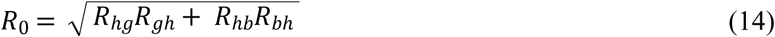

For vector-borne viral transmission between two humans, two stages of the transmission process are involved: the transmission from human “A” to mosquito “B” (generation 1), and then from mosquito “B” to another human “C” (generation 2). The number of mosquitoes “B” caused by an infectious human “A” is R_bh_ (or R_gh_), and the number of humans “C” caused by each infectious mosquito “B” is R_hb_ (or R_hg_). After two generations, the total number of secondary human-to-human cases for both mosquito species is R_hg_R_gh_ + R_hb_R_bh_. Therefore, the basic reproductive number (R_0_), which characterizes the number of cases in one generation, is the geometric average of the cases in two generations, that is 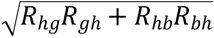.

### Sensitivity analysis

Because the transmission parameters are only known approximately, it is important to understand how variations in these parameters affect model outcomes. To quantify the impact of changes in parameters on R_0_, three types of sensitivity analysis were performed: a local sensitivity analysis, an extended sensitivity analysis, and a global sensitivity analysis.

In the local sensitivity analysis, sensitivity indices were derived to quantify how small changes in the parameter of interest *p* caused variability in the model output of interest *q.* If an input parameter *p* changed by *x%*, then the output quantity *q* changed by 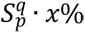. As such, the sensitivity index’s magnitude determines the relative importance of the model parameters on the model predictions. The sign of the sensitivity index indicates the direction of change of the output in response to the parameter change. The sensitivity indices of R_0_ were analytically computed by evaluating partial derivatives of R_0_ (Eq. 9) with respect to each parameter of interest at the baseline value, multiplied by a scaling factor 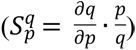. As a result, the local sensitivity indices are valid only at a small range around the parameter baseline values.

In the extended sensitivity analysis, the responses of R_0_ to the variations in each parameter of interest are calculated over the entire possible range of that parameter (Table 1), while fixing all other parameters at their baseline. The extended local sensitivity analysis curves were plotted to depict the derivative of R_0_ as a function of the model parameter of interest at all values within its possible range.

In the global sensitivity quantification, the values of R_0_ were calculated using multiple combinations over the full range of all the parameters. The parameters were treated as random variables (all parameters can simultaneously take any values within their possible ranges), and R_0_ had a distribution, which depended on the distributions of parameters. In this analysis, each of the model parameters was assumed to vary independently from each other and has a uniform distribution. The description of sensitivity analyses was given in more detail in the previous publication [20]. All sensitivity analyses were done in MATLAB R2018a (version 9.4.0).

## Results

### Exploratory survey of mosquito bite exposure in adults in the Greater New Orleans Region

In the preliminary survey, the total number of retrieved questionnaires was 104 (ORL, 33; JEF, 24; TAM, 47). The average return rate across study sites was 20.7%. The results are shown in Fig 1

**Fig 1.**
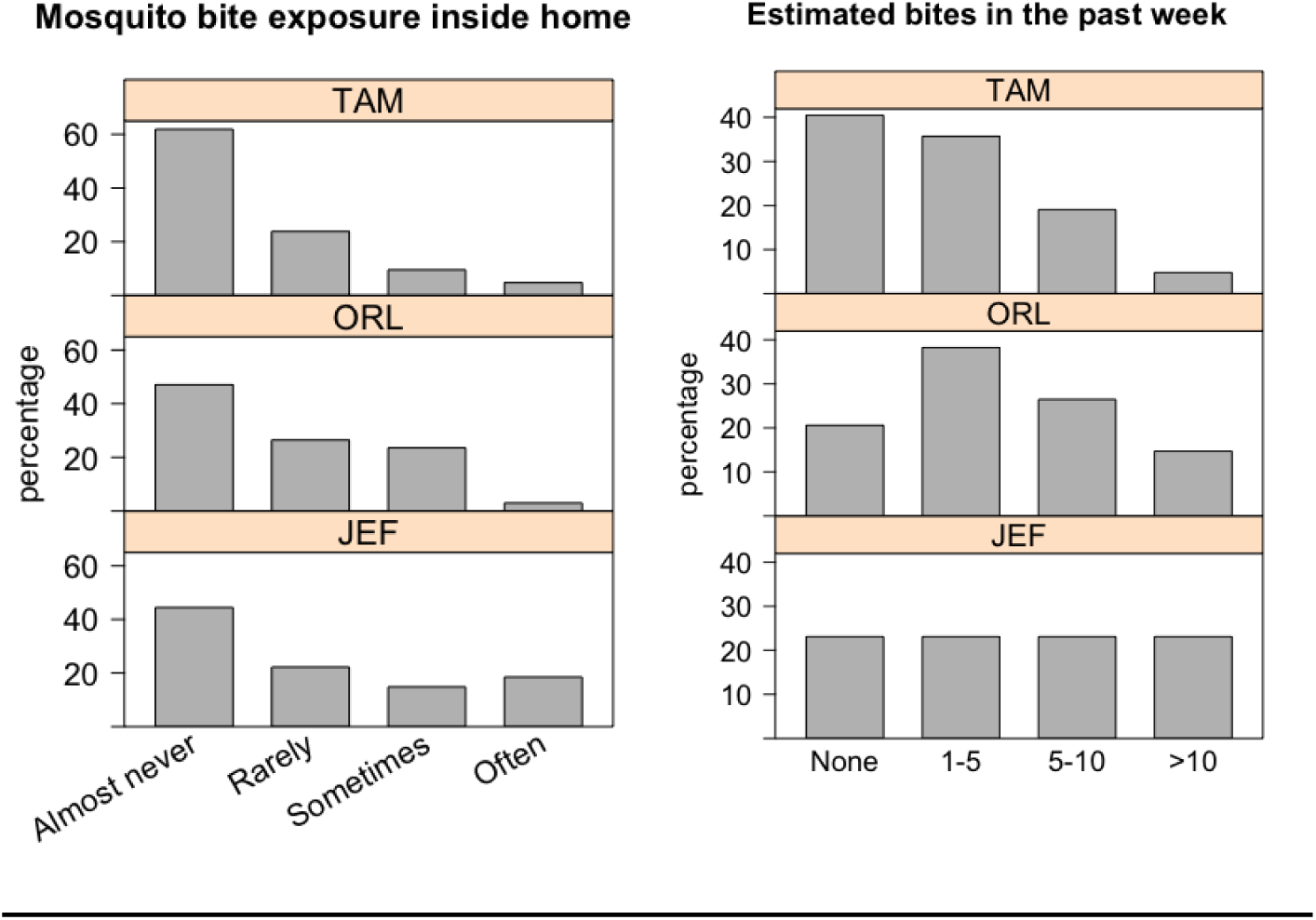
Results from the preliminary survey showing the frequency of bite exposure inside homes and the estimated numbers of bites research participants experienced in the past 7 days. The percentages of participants choosing each answer of the multiple-choices questions are shown.

The preliminary results suggested variations between study sites. Research participants in JEF reported higher exposure to mosquito bites than research participants in ORL and TAM. In TAM, around 40% of research participants indicated that they did not receive any mosquito bites in the past 7 days. While in ORL, 38% of research participants chose “1-5” bites in the past 7 days. In JEF, equal proportions (23%) of research participants reported being bitten more than 10 times, 5-10 times, 1-5 times, and none in the past 7 days.

When asked how often they experienced mosquito bites inside of their homes, 19% of research participants from JEF chose “often” as the answer, higher than the other two study sites (both were <5%). In all study sites, the place where people most often experienced outdoor mosquito bites was around their homes (78%, 72%, 56% for TAM, JEF, and ORL, respectively). In ORL, “public space” was also reported as a place where people most often experienced mosquito bites (32%).

### Mosquito bite exposure rates in adults in the Greater New Orleans Region

For the second survey, a total of 941 and 801 questionnaires were distributed in ORL and TAM, respectively. The average numbers of addresses per block were 23.53 (SD = 7.90) for ORL and 20.03 (SD = 3.44) for TAM. In ORL, a total of 91 questionnaires were retrieved, with an average return rate of 10.06% (SD = 6.46%) per block. In TAM, a total of 94 questionnaires were retrieved, with an average return rate of 11.35% (SD = 8.26%) per block.

The average numbers of adults (>18 years old) per household were 1.84 for ORL (SD = 0.73) and 2.11 for TAM (SD = 0.62). Graphs showing the gender and age distribution of research participants in both study sites are shown in Supplementary Figure 2. In total, research participants included 90 females, 70 males, and 25 individuals who did not indicate their gender. Of these, one person was between 18-25 years old, 38 were between 26-40 years old, 78 were between 41-65 years old, 63 were more than 65 years old, and 5 failed to indicate their age range.

Overall, the reported numbers of mosquito bites that occurred outdoors and indoors within the past 24 hours in ORL, after adjustment with sampling weights, were 5.48 (SE=0.90) and 1.68 (SE=0.46), respectively. The reported numbers of bites that occurred outdoors and indoors within the past 24 hours in TAM, after adjustment with sampling weights, were 2.28 (SE=0.34) and 0.32 (SE= 0.13), respectively. In both study sites, the average numbers of reported bites that occurred outdoors were significantly higher than indoors (Wilcoxon rank sum test, ORL: df = 34, p-value <0.001, TAM: df = 30, p-value <0.001). In addition, the reported numbers of bites were significantly higher in ORL compared to TAM for both outdoor and indoor settings (Wilcoxon Rank Sum test, outdoors: df = 66, p-value= 0.003; indoors: df = 66, p-value <0.001). The average reported numbers of bites that occurred outdoors and indoors, after adjustment with sampling weights, within the past 24 hours in both study sites across months are shown in Fig 2.

**Fig 2.**
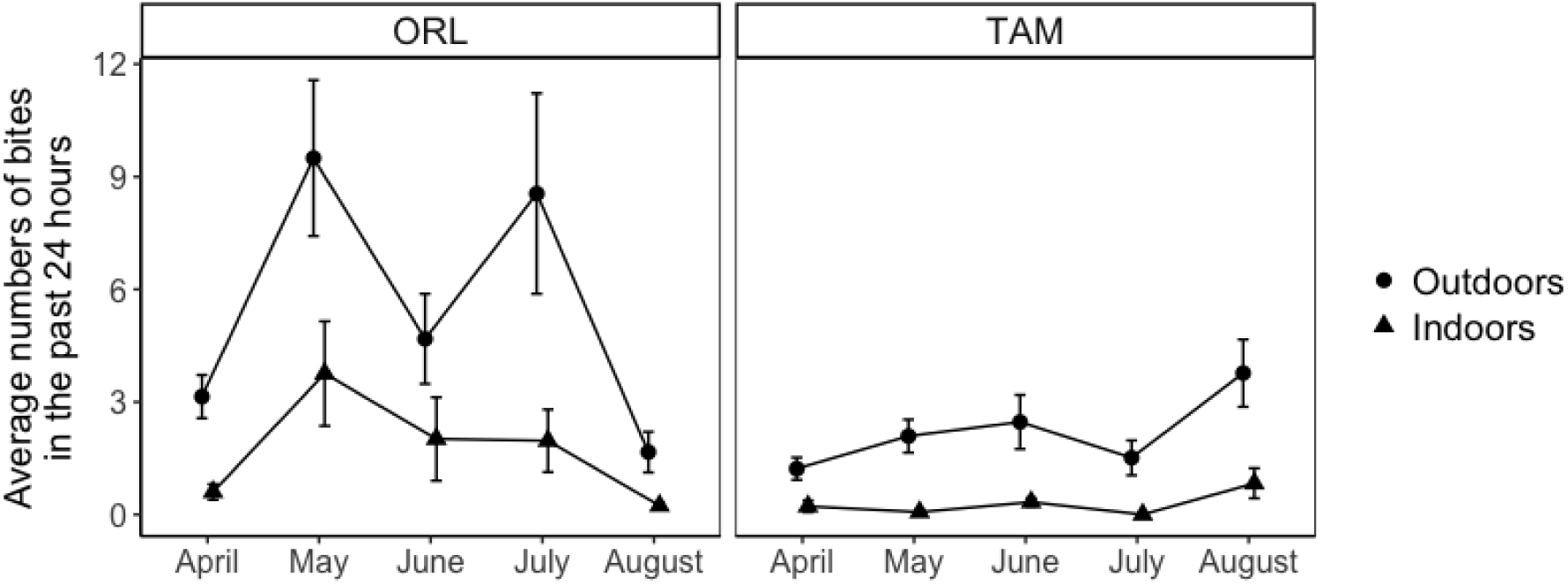
The average numbers of mosquito bites per person, after adjustment with sampling weights, in the past 24 hours that research participants reported are shown by sites and month of data collection. The circles represent the outdoor bites and the triangles represent indoor bites. Error bars represent the standard errors.

### Factors affecting bite exposure in adults in the Greater New Orleans Region

For research participants who reported receiving outdoor mosquito bites within the past 24 hours, they were asked to indicate the locations that they experienced these bites. In TAM, 47 participants or around 90% reported being bitten around their homes (answers such as ‘front yard’, ‘backyard’, ‘on dock’, ‘sitting in my open garage’), whereas 5 participants or around 10% reported being bitten both around their homes *and* at public spaces (answers such as ‘backyard and dog park’ and ‘yard and during a walk’). In ORL, 33 participants or around 59% reported being bitten around their homes (answers such as ‘backyard’, ‘front porch’, and ‘side yard’), 6 participants or 11% reported being bitten at public spaces (answers such as ‘outside while at work’, ‘while walking the dog’, and ‘walking along Crescent City park and inside of an indoor/outdoor bar’), and 17 participants or 30% reported being bitten both around their homes *and* at public spaces (answers such as ‘backyard, while out walking’ and ‘Clouet garden and my backyard’).

Information about the time spent outdoors within the past 24 hours was collected from research participants (Supplementary Figure 3). After adjustment with sampling weights, research participants in ORL spent 41.78 minutes (SE = 7.43) and 67.47 minutes (SE = 6.91) outdoor during the weekday and weekend on average, respectively. After adjustment with sampling weights, research participants in TAM spent 54.33 minutes (SE = 5.53) and 51.00 minutes (SE = 8.17) outside during the weekday and weekend on average, respectively. The difference of the time spent outside between the weekend and weekday was significant for research participants in ORL (Wilcoxon Rank Sum test, df = 34, p-value = 0.02) but not for research participants in TAM (Wilcoxon Rank Sum test, df = 30, p-value = 0.3). In addition, the difference of the time spent outside between research participants in ORL and TAM was statistically significant for the weekend (Wilcoxon Rank Sum test, df = 32, p-value = 0.02) but not during the weekday (Wilcoxon Rank Sum test, df = 32, p-value = 0.3).

The time spent outside during the time period between 5 pm to 8 pm (or evening time), and 8 pm to 6 am (or nighttime) showed significant correlations with reported bite numbers using Spearman’s correlation test. The correlation coefficient was 0.25 (p-value = 0.003) and 0.28 (p-value <0.001) for the evening and nighttime, respectively. The time spent outside during the time period between 6 am to 10 am (or morning time), and 10 am to 5 pm (or daytime) did not show significant correlations with reported bite numbers (Spearman’s correlation test; p-value = 0.078 and 0.975, respectively).

A generalized linear model analysis was used to determine which variables are associated with how much time the research participants reported spending outside in the evenings and at night. A table showing the model’s result is shown in Supplementary Table 2. Only the age range of research participants and the weekend/weekday setting showed significant associations with the time participants reported spending outside in the evening and night. Specifically, older participants spent less time outside in the evening and night than younger participants. Research participants also spent less time outside on weekdays than on weekends.

Another generalized linear model analysis was used to determine the effect of study site, the month of data collection, total time spent outside in the evening and night, and gender of research participants on the reported numbers of outdoor bites. The results, detailed in Supplementary Table 3, indicated that the time spent outside in the evening and night, the month of data collection (May, July, and August), and study site show significant associations with the reported outdoor bite numbers. The results show that, when controlled for other variables including the time they spent outside, research participants in ORL reported experiencing higher mosquito bites than participants from TAM. Gender did not show a significant association with the reported bite numbers (p-value = 0.053).

### Determining mosquito species contributing to bite exposure in the Greater New Orleans using Human Landing Capture

The average composition of female mosquito species and types captured during HLC in both study sites are shown in the top graphs of Fig 3. In ORL, on average 56.02% of landed female mosquitoes were *Ae. albopictus* and 36.13% were *Ae. aegypti*. In TAM, on average 50.98% of landed mosquitoes were *Ae. albopictus* and 28.43% were *Ae. aegypti*. In ORL, species other than *Ae. aegypti* and *Ae. albopictus* that were captured included: *Ae. taeniorhynchus*, *Ae. vexans*, *Mansonia titillans* and *Ae. infirmatus*. In TAM, other species included: *An. bradleyi*, *Cx. salinarius*, *Cx. restuans*, *Ae. taeniorhynchus*, and *Ae. sollicitans*.

**Fig 3.**
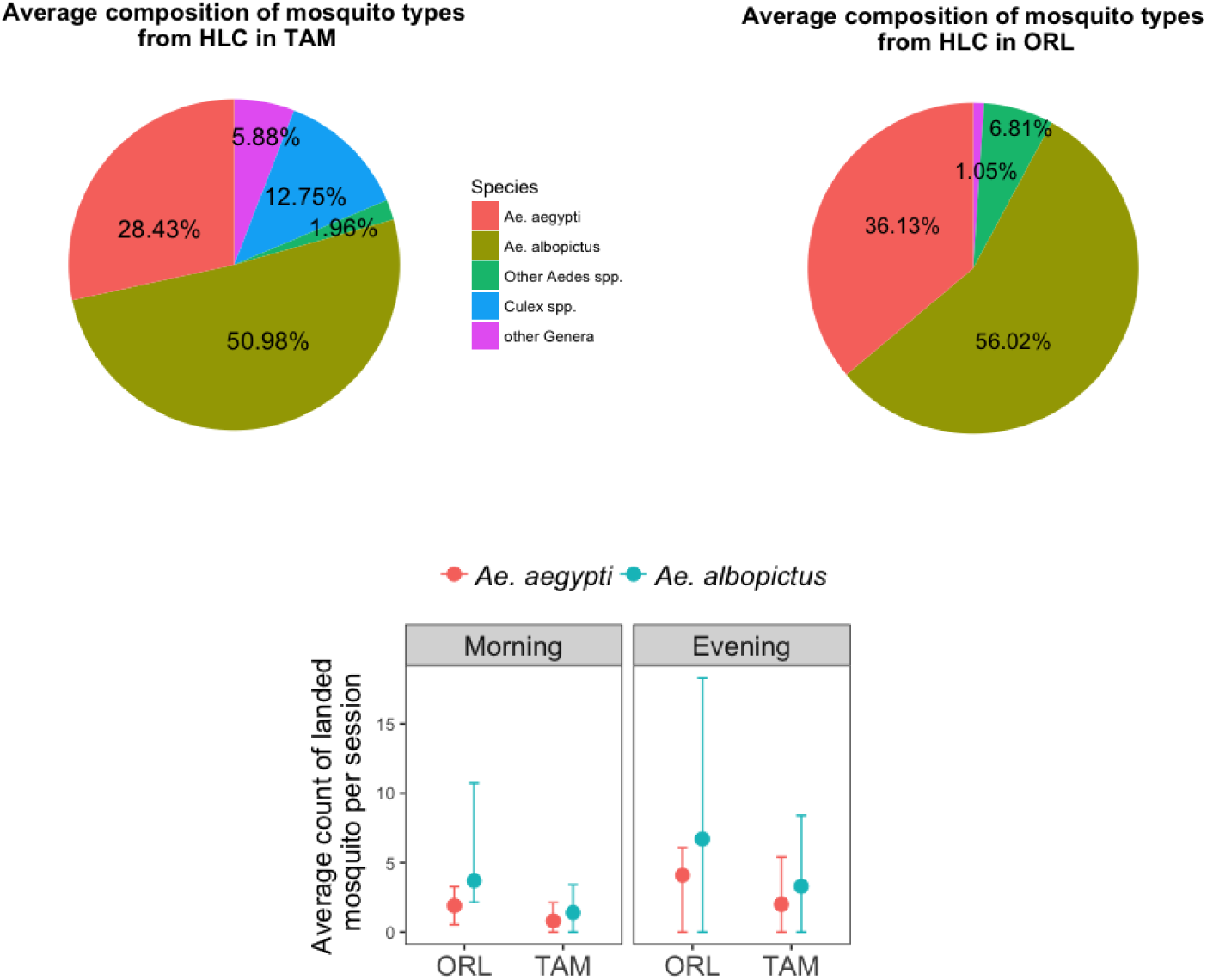
Top: pie graphs showing the average composition of mosquito types captured during HLC in TAM and ORL. Bottom: average numbers of landed female *Ae. aegypti* and *Ae. albopictus* in ORL and TAM during the 1.5 hour of HLC sessions in the morning and evening.

The average numbers of female *Ae. aegypti* and *Ae. albopictus* landed during 40 HLC sessions are shown in the bottom graph of Fig 3. In ORL, the average numbers of landed female *Ae. aegypti* in the morning and evening HLC session (1.5 hour) were 1.9 (SD = 1.37) and 4.1 (SD = 1.97), respectively. The average numbers of landed female *Ae. albopictus* in the morning and evening HLC session were 3.7 (SD = 7.03) and 6.7 (SD = 11.60), respectively. In TAM, the average numbers of landed female *Ae. aegypti* in the morning and evening HLC session were 0.8 (SD = 1.32) and 2.0 (SD = 3.40), respectively. The average numbers of landed female *Ae. albopictus* in the morning and evening HLC session were 1.4 (SD = 2.01) and 3.3 (SD = 5.10), respectively. Averaging data from both study sites, the number of landed mosquitoes was higher in the evening than in the morning for both *Aedes* species. However, the difference is statistically significant only for *Ae. aegypti* and not for *Ae. albopictus* (Wilcoxon Rank Sum test, p-value = 0.04 and 0.08, respectively). In addition, averaging data from both morning and evening sessions, the number of landed mosquitoes in ORL was significantly higher than in TAM for *Ae. aegypti* but not for *Ae. albopictus* (Wilcoxon Rank Sum test, p-value = 0.002 and 0.2, respectively).

### Basic Reproductive Number (R_0_) and the initial transmission of DENV

The model analysis simulated a situation where one infectious human was introduced into fully susceptible populations of humans and mosquitoes. Table 2 shows the result from the model analysis using different values of local human-mosquito contact rates, calculated using equation (7) and (8), while holding other parameters at baseline values. The output of interest includes R_0_, the percentage of infected and recovered human at their peaks, and the numbers of days before the number of infected and recovered human reach their peaks.

**Table 2.** Results from the model analysis using different values of local human-mosquito contact rates.

| Parameter | R<sub>0</sub> | Infected human at its peak |  | Recovered human at its peak |  |
| --- | --- | --- | --- | --- | --- |
|  |  | Percentage over total population | Time at the peak (day) | Percentage over total population | Time at the peak (day) |
| Using human-mosquito contact rates from ORL |  |  |  |  |  |
| baseline values | 2.41 | 6.53% | 188 | 97.00% | 332 |
| minimum values | 1.56 | 1.30% | 510 | 64.70% | 907 |
| maximum values | 3.26 | 10.83% | 124 | 99.67% | 213 |
| Using human-mosquito contact rates from TAM |  |  |  |  |  |
| baseline values | 0.73 | - | - | - | - |
| minimum values | 0.50 | - | - | - | - |
| maximum values | 0.96 | - | - | - | - |

Given values of human-mosquito contact rates acquired from both study sites, only the R_0_ during the initial DENV transmission in ORL exceed 1. When using the baseline value of human-mosquito contact rate from ORL, the calculated R_0_ for DENV transmission in the area was 2.41 and the infected human number peaked at day 188^th^ after the virus introduction. When using the minimum value for the contact rate from ORL, R_0_ was greater than 1 even though the outbreak was less explosive. The infected human number peaked at day 510^th^ after the initial virus introduction. R_0_ value was highest (3.26) for the maximum value of the contact rate from ORL, and the number of infected humans peaked at day 124^th^. However, none of the human-mosquito contact rate values quantified in TAM resulted in an R_0_ exceeding 1, and therefore a small initial infection would die out.

Given the baseline value of human-mosquito contact rate in ORL, the number of infected *Ae. aegypti* at its peak was 4,647. This is higher than infected *Ae. albopictus*, where their number at the peak was 182 (Fig 4). When using the maximum value of human-mosquito contact rate in ORL, the number of infected *Ae. aegypti* and *Ae. albopictus* at their peaks were 8,779 and 360, respectively. Finally, when using the minimum value of human-mosquito contact rate in ORL, the number of infected *Ae. aegypti* and *Ae. albopictus* at their peaks were 713 and 27, respectively.

**Fig 4.**
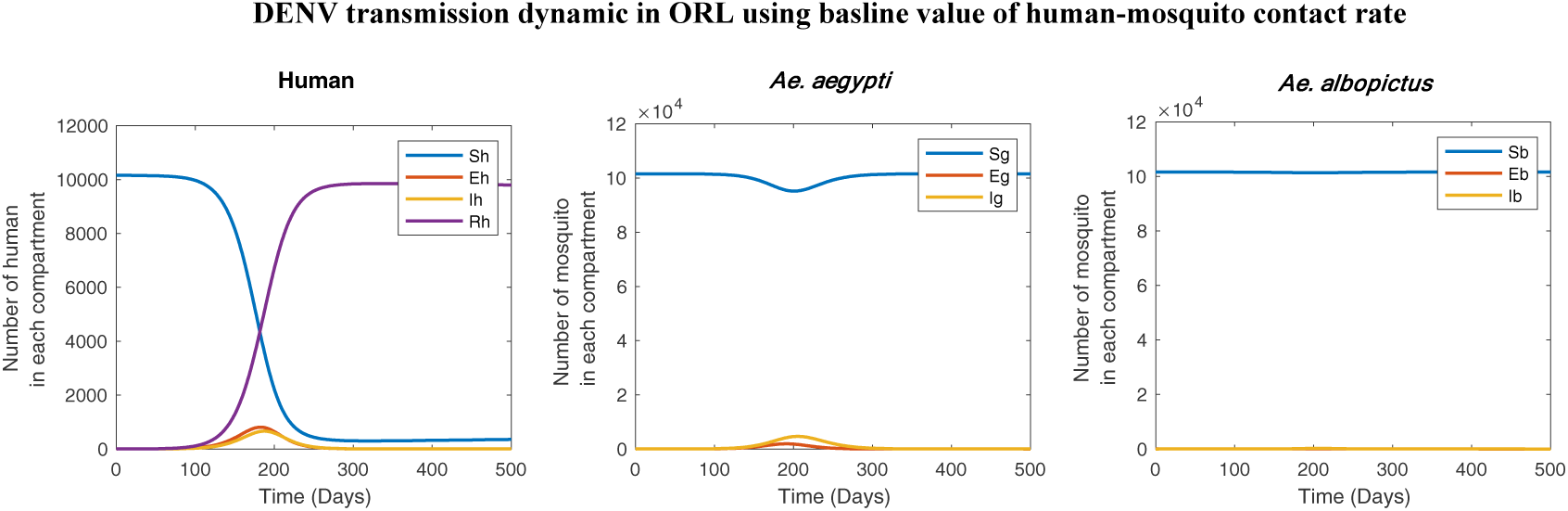
Model analysis of DENV transmission in ORL using baseline value of human-mosquito contact rate.

### Local sensitivity analysis

The local sensitivity indices of R_0_ with respect to model parameters are shown in Table 3. For both transmission scenarios in ORL and TAM, the R_0_ is most sensitive to 1) *Ae. aegypti*-human contact rate (B_g_), 2) the probability of DENV transmission from mosquito to human given an infectious bite (β_h_), and 3) the recovery rate of human (or the inverse of viremic period; γ_h_), evaluated at their baseline values. At the baseline values, the basic reproductive number is least sensitive to the inverse of the intrinsic incubation period (ν_h_) and human death rate (μ_h_).

**Table 3.**
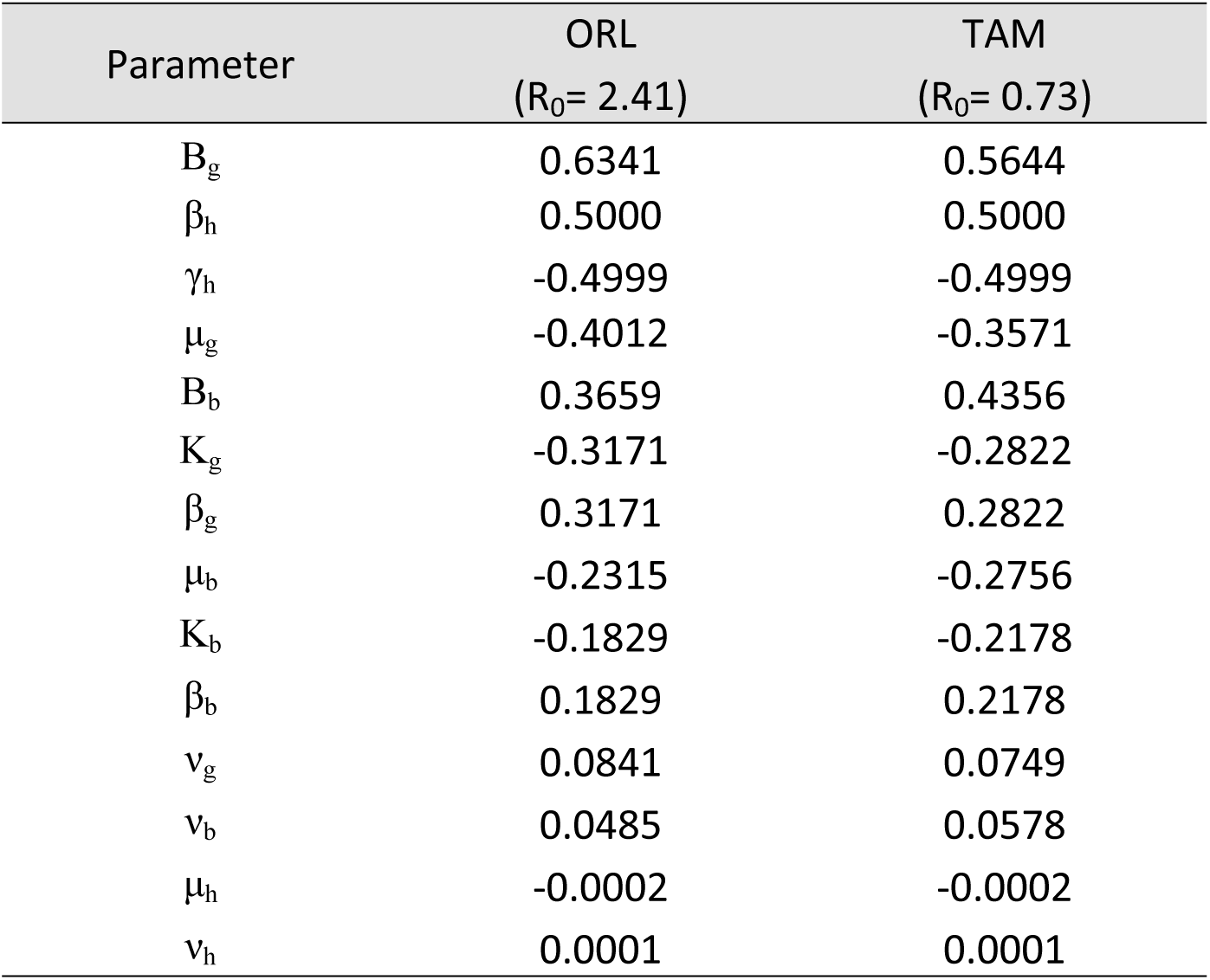
Sensitivity indices of R_0_ with respect to model parameters at the baseline values.

| Parameter | ORL<br>($R_0 = 2.41$) | TAM<br>($R_0 = 0.73$) |
| --- | --- | --- |
| $B_g$ | 0.6341 | 0.5644 |
| $\beta_h$ | 0.5000 | 0.5000 |
| $\gamma_h$ | -0.4999 | -0.4999 |
| $\mu_g$ | -0.4012 | -0.3571 |
| $B_b$ | 0.3659 | 0.4356 |
| $K_g$ | -0.3171 | -0.2822 |
| $\beta_g$ | 0.3171 | 0.2822 |
| $\mu_b$ | -0.2315 | -0.2756 |
| $K_b$ | -0.1829 | -0.2178 |
| $\beta_b$ | 0.1829 | 0.2178 |
| $v_g$ | 0.0841 | 0.0749 |
| $v_b$ | 0.0485 | 0.0578 |
| $\mu_h$ | -0.0002 | -0.0002 |
| $v_h$ | 0.0001 | 0.0001 |

The sign of the sensitivity index indicates the relationship between the direction of changes in R_0_ and model parameters. For example, the sensitivity indices of R_0_ with respect to human-mosquito contact rates (both B_g_ and B_b_), evaluated at their baseline values, are positive. Therefore, as the contact rate between mosquito and human increases, the R_0_ also increases. On the contrary, the sensitivity indices of R_0_ with respect to γ_h_, evaluated at their baseline values, are negative. As a result, as the human recovery rate increases (i.e. viremic period decreases), the R_0_ decreases. Another observation is the negative value of the sensitivity indices of R_0_ with respect to the mosquito carrying capacity (both K_g_ and K_b_), evaluated at their baseline values. This can be interpreted that as the mosquito carrying capacity increases, the R_0_ decreases. The mathematical explanation for this unexpected relationship is discussed in the Discussion section.

The relative ranking of the parameter importance was almost the same between the two scenarios (Table 3). The only exception is that B_b_, or *Ae. albopictus*-human contact rate, becomes relatively less important at determining R_0_ in the ORL scenario compared to TAM. This results from the assumption that *Ae. Albopictus* has a lower vector competence than *Ae. aegypti*, and *Ae. aegypti*-human has a higher contact rate in the ORL.

### Extended sensitivity analysis

The extended sensitivity analysis plots of R_0_ with respect to the mosquito-human contact rate for the transmission scenario in ORL are shown in Figure 6. The extended sensitivity analysis plots of R_0_ to other selected model parameters for ORL and TAM are shown in Supplementary Figure 4 and 5, respectively.

First, consider the top two graphs of Fig 5, which show how the R_0_ value changes in response to changes in the *Ae. aegypti*-human contact rate (B_g_; top left panel) and the *Ae. albopictus*-human contact rate (B_b_; top right panel), while holding all other parameters at their baseline values. Both plots show curves with positive trends, indicating that a decrease in contact rate, while holding other parameters at their baselines, will cause R_0_ to decrease. However, this relationship is not linear; as the contact rate decreases, the slope becomes smaller. That is, the reduction in human-mosquito contact rate, when focused on only one vector species at a time, becomes less effective at reducing R_0_ when the contact rate is already small. In fact, in the ORL scenario, reducing the contact rate between humans and only one vector species at a time will fail to reduce R_0_ below 1. This is because the contact rate between humans and the other vector species is high enough to maintain the transmission.

**Fig 5.**
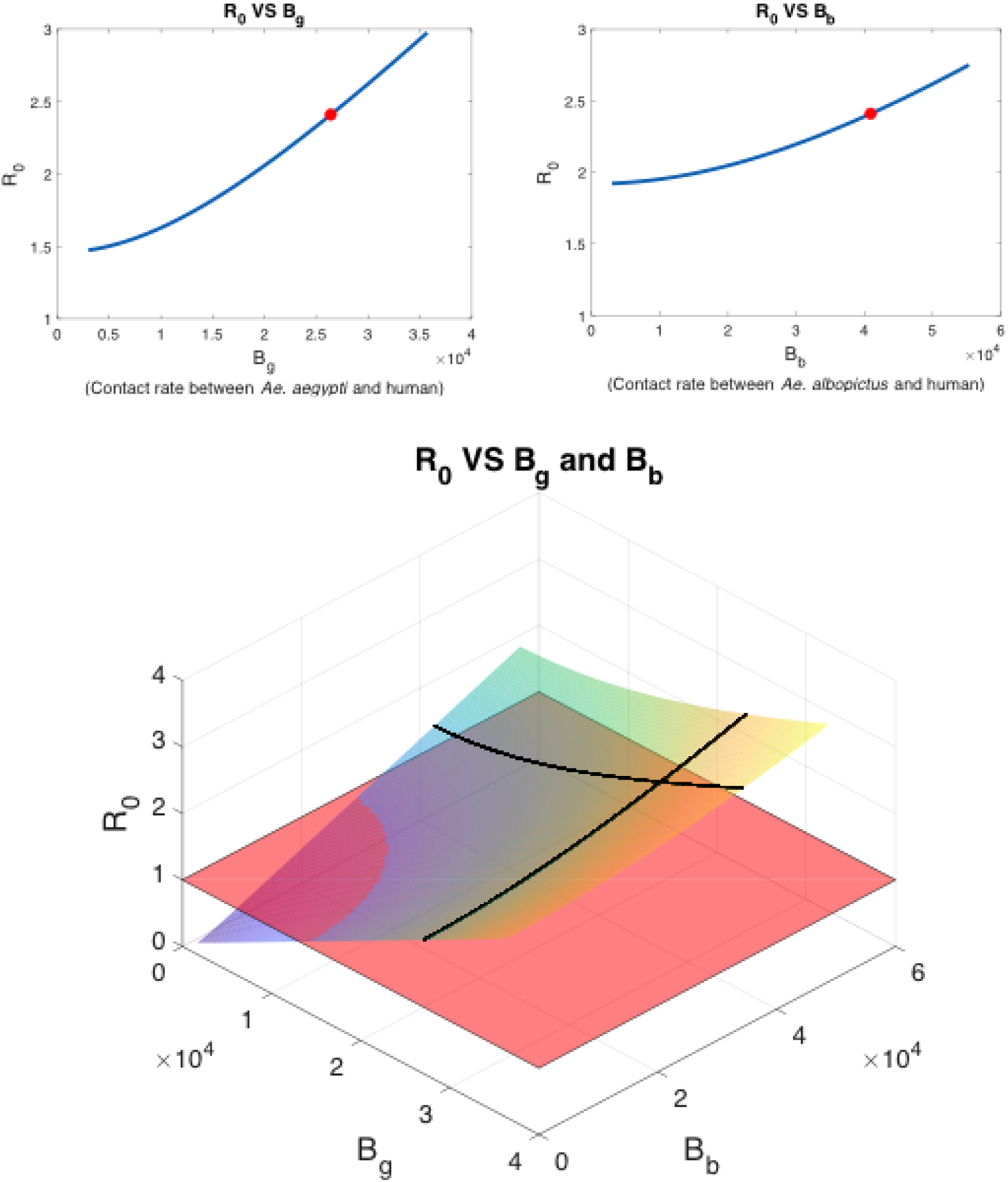
The extended sensitivity analysis plots of R_0_ with respect to the human-mosquito contact rate for the model analysis of DENV transmission in ORL. In the top graphs, the red dots represent the R_0_ at the contact rate baseline values. In the bottom graph, the red plane represents where R_0_ = 1 and the black lines represent the R_0_ values at baseline contact rates of each of the two *Aedes* species and humans. The point where the two lines meet represents the R_0_ value at the baseline contact rates of both *Aedes* species and humans.

Next, consider the bottom graph in Fig 5, which shows how R_0_ changes in response to the changes in both B_g_ and B_b_ simultaneously, while holding other parameters at their baseline values. In this case, the reduction of both B_g_ and B_b_ at the same time below certain threshold values will result in R_0_ < 1.

### Global sensitivity analysis

Figure 7 shows the distribution of R_0_ calculated from combinations of model parameter values, which were sampled uniformly and independently within their possible ranges. The R_0_ distribution for the ORL scenario was wider at the base and had a longer tailed distribution, indicating that there was a higher variation in the outcomes. The percentage of scenarios (or the combinations of parameter values) that resulted in an R_0_ > 1 indicated how likely DENV was to spread in either location. In the ORL case, 74.52% of scenarios resulted in an R0>1. In TAM, 68.80% of scenarios resulted in an R0>1. As such, ORL was more receptive to an initial outbreak of DENV than TAM.

**Figure 7.**
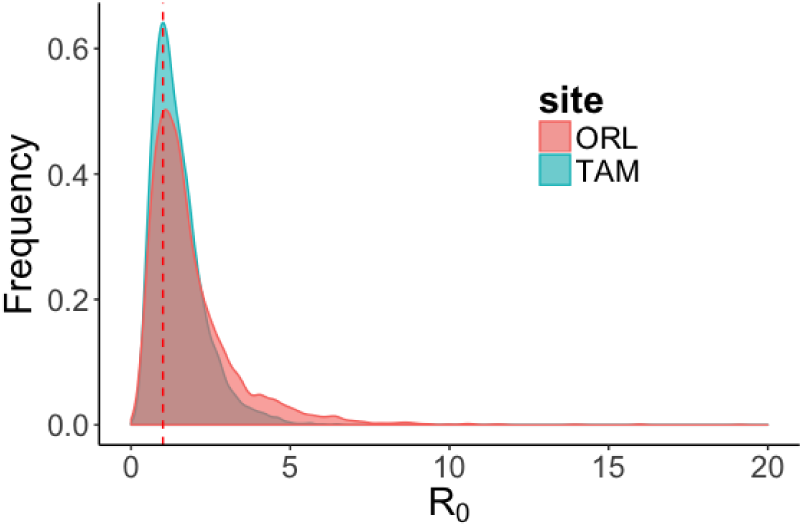
The frequency distribution of R_0_ values calculated from combinations of model parameter values sampling uniformly and independently. The vertical dashed line at R_0_=1 indicates the threshold value for an outbreak.

## Discussion

Mosquito bite exposure was investigated using a questionnaire survey to ask research participants about their past experience receiving mosquito bites. We found that the mosquito bite exposure on research participants occurred more frequently in the outdoors than indoors in both study sites. The location that research participants most often reported being exposed to mosquito bites was around their homes. We quantified the correlation between the reported bite number and the time spent outside in the evenings and at night. After controlling for the time duration spent outside, there was a significant effect of study site on the outdoor biting rate, where participants in ORL reported receiving more mosquito bites than participants in TAM. In places such as the Greater New Orleans Region where the mosquito bite exposure between indoors and outdoors may be different, the human-mosquito contact rate depends on the density of host-seeking female mosquitoes and human behavior, such as the time spent outside.

Interestingly, the indoor bite exposure rate was also higher for ORL than in TAM. The potential reason for this difference was not investigated in this study. According to the 2016 ACS 5-year estimates, the median household income in TAM is 42% higher than in ORL (Supplementary Table 1). It is possible that factors such as the integrity of the wall, the availability of air conditioners, combined with human behaviors (keeping doors or windows open) determine the difference in indoor bite exposure rate [37]. Future study is needed to investigate the relative importance of these factors on indoor mosquito bite exposure.

Only a few other studies have used surveys to investigate mosquito bite exposure. A study by Dowling *et al.* asked research participants in suburbs of Washington DC how often they were bitten by mosquitoes. Out of 246 participants, 48% chose ‘Everyday’, 28% chose ‘Few days a week’, and 24% chose ‘Few days a month or fewer’ [38]. A similar study by Halasa *et al.* interviewed residents in two counties of New Jersey and found that during a typical summer week, 80.2% of respondents reported being bitten at least once and 77.7% were bitten while outdoors [39]. In Halasa’s study, bite exposure occurred most often in the evening (52.1%), followed by at night (31.4%), and late afternoon (30.6%). A study by Read *et al.* used a unique study design to compare the number of mosquito bites that participants thought they received while sitting outside for 5 minutes with the number of mosquitoes captured concurrently on a staff person using a Whole Person Bag Sampler [40]. The study showed that respondents’ reported bites received during the 5-min blinded test time increased with increasing trap count. However, there was a higher discrepancy between the reported bites and the trap count at the lower trap count.

The HLC data from this study indicated that there were higher numbers of host-seeking mosquitoes in ORL than in TAM, and more in the evening than in the morning. Even though this study was not designed to compare the bite survey to HLC, the observations from both methods were congruous. For example, the higher reported mosquito bite exposure in ORL mirrored the higher number of host-seeking mosquitoes in that site, compared to TAM. In addition, the correlation between the reported outdoors time and the amount of mosquito bites was found only in the evening and nighttime, but not in the morning. This finding was consistent with our HLC data and other studies, which found higher numbers of host-seeking *Ae. aegypti* in the evenings than in the mornings [24, 41]. Future study is needed to investigate the correlation between the reported bite exposure level from surveys and the number of landed mosquitoes from HLC experiments.

Our model analysis showed that the human-mosquito contact rate played an important role in determining contrasting outcomes in dengue transmission simulated in the two study sites. The local sensitivity indices indicated that the contact rate between humans and *Ae. aegypti* was the most important parameter determining the R_0_, and was more important that the contact rate between humans and *Ae. albopictus*. This was because of the difference in the vector competence between the two species. *Ae. aegypti* is thought to be a more competent vector [42] and we set its vector competence value to be higher. Our laboratory experiment to test vector competence of the locally collected mosquitoes also suggested that the local *Ae. aegypti* was more competent than the local *Ae. albopictus* (data not shown).

Interestingly, changes in the carrying capacity of mosquitoes (which controlled their population size) showed an inverse relationship with the changes in R_0_, while holding other parameters at their baselines. That is, as the mosquito population size decreases, then the potential for disease outbreak increases. This is counter-intuitive because one may expect the risk of an outbreak to be smaller when the vector density is low. However, the assumption of this model is that the contact rate is frequency-dependent: it does *not* depend on human or mosquito density.

This assumption may be valid when human, and mosquito variables contribute to a fixed amount of bites that is compromised by both the mosquito’s desire to blood-feed and the number of bites humans can tolerate. Under this assumption, the biting rate *per mosquito* (B_g_/K_g_ and B_b_/K_b_) would increase as the carrying capacity (K_g_ and K_b_) of mosquitoes decreases. The increase in this biting rate *per mosquito* results in a higher outbreak potential. Even though this explanation is justifiable mathematically, the real-world mechanisms will likely be more complicated, and may result in a different transmission outcome. Nonetheless, when designing a mosquito-borne disease control program, especially in endemic areas, control tools that reduce contact between human and mosquito should be implemented along with those that reduce mosquito density.

Mathematical models are a simplified simulation of a real world complex process. As such, the models are biased and limited by their assumptions and parameter values. In our model, we assumed uniform distributions of human and mosquito density in both space and time. In reality, this is unlikely to hold true. For example, the mosquito population size in the Southern US fluctuates significantly as a response to seasons. When the simulated time period spans across several seasons, then the model parameters need to account for the fluctuating mosquito’s carrying capacity and death rate.

In addition, a deterministic model was utilized. Even though this model type has been applied in many disease systems due to its simplicity and clarity [43, 44], it ignores heterogeneity and stochasticity inherent in natural disease transmission. Early in the disease invasion stage, when there are only a few infectious hosts, stochasticity and chance events often play an important role in determining the transmission course [45]. For example, infectious hosts can all heal or die due to chance alone before transmission can take off even when R_0_ is above one.

We also assumed that the contacts were evenly distributed among individuals. This assumption rarely applies in the real world. Often, only a small fraction of individuals, known as super-spreaders, contribute significantly to contacts and transmission events [46]. Studies have shown that mosquito biting and bite exposure are associated with many variables such as human body size, alcohol consumption, skin odor, housing type, or proximity to mosquito habitats [47]. In addition, behavioral changes that may be associated with more severe human cases (e.g. house-ridden individuals) could result in differential bite exposure rates. Questionnaire-based surveys may be a valuable tool that could be feasibly used to investigate how these factors impact heterogeneity in mosquito bite exposure among individuals.

Another important factor determining the accuracy of the model’s predictions is the accuracy of the parameters’ values. Human-mosquito contact rate has rarely been characterized in the field and is among the least known parameters in mosquito-borne disease transmission. HLC has been the traditional gold standard method, but its use is often impractical [48] and does not take into account human lifestyles or other innate human variables. Molecular approaches to profile the mosquito blood meal are expensive, time-consuming, and can only provide biting patterns [49–51] and not rates (but see [52]). The use of questionnaire-based surveys, especially in the form of door-hanger questionnaires, provides a low-cost, fast, and feasible alternative.

Despite their benefits, using surveys to approximate human-mosquito contact rates may result in some biases. For example, in an attempt to get a full blood meal, a mosquito may probe repeatedly on a host [53]. As a result, a person may report being bitten multiple times but the contacts were with only one mosquito. In additions, the bites research participants received could be from arthropods other than mosquitoes. Even though the participants were asked to indicate the number of mosquito bites within the past 24 hours (instead of the past 7 days, as was done in the preliminary survey), it is likely that there was a recall bias. To reduce this bias, a prospective cohort study design could be used in future studies. In addition, only a small portion (∼10%) of the targeted population participated in the study. This may cause selection bias because the decision to participate in the study may reflect inherent characteristics of the participants. Subjects who decided to take part in a survey may have a strong interest or awareness in the study topic [54]. By using other sampling methods or increasing sample size, selection bias could be reduced.

Another limitation in our study results from the use of a small-scaled HLC to characterize the mosquito compositions only at crepuscular periods. The composition of mosquitoes that may contribute to bites during nighttime was not characterized. We expected that nighttime biters such as *Culex* spp. and *Anopheles* spp. may contribute considerably to bites during this period.

Computational uncertainties are unavoidable in predicting the dynamics of an epidemic. The baseline model parameters in Table 1, together with the human-mosquito contact rates obtained through the survey, are only our best-guess estimates of the model parameters. Such uncertainties in the parameters could affect the reliability of the model predictions. It is important to emphasize that the quantitative values of the model outputs, such as R_0_, should not be taken at face value. They only give us insight into potential outcomes of disease spreads. Fortunately, the qualitative aspects of the model, such as the relative importance of the different factors are usually robust and less sensitive to these assumptions.

The probability of a disease emergence in a new geographical area encompasses two qualitative attributes: vulnerability and receptivity [55]. Vulnerability indicates the influx of infected individuals into an area of interest, while receptivity reflects the local conditions that are conducive for disease transmission. In this study, the risk of DENV outbreak was investigated only at the level of receptivity. In Louisiana, a total of 45 imported cases were reported from 1980 to 2015 (Dengue Annual Report, Louisiana Office of Public Health, 2015). In general, despite the highly receptive condition, the probability of a DENV outbreak could be lower due to its low vulnerability.

In conclusion, we found that the use of a questionnaire-based survey is a feasible method to estimate human-mosquito contact rates. It can be used to compare mosquito bite exposure levels between settings in order to evaluate how environmental factors and intervention strategies may impact disease risk. Most importantly, it may provide an avenue to investigate how changes in human characteristics such as behaviors, lifestyles, use of clothing and personal protection, and other innate variables affect mosquito bite exposure and the risk of infection in a way that is very difficult to do with HLC. This information is indispensable if we want to predict how the changing environment due to unplanned urbanization, poverty, and climate change impacts mosquito-borne disease transmission. In addition, the use of mathematical models to simulate disease transmission produces valuable information that helps us understand how changes in the transmission variables may impact disease transmission. This type of knowledge facilitates the planning of cost-effective disease prevention programs to target the most important transmission factor which may lead to the largest reduction in transmission risk.

## Author contribution

PT participated in conceptualization and study design, conducted field research, curated data, participated in mathematical model development, and performed formal analysis and data visualization. ZQ participated in model development and analysis, and participated in data visualization. JOY participated in study design and data analysis, and validated mathematical and statistical analyses. JMH participated in mathematical model development, and validated mathematical model analyses. DMW participated in conceptualization and study design, oversaw and coordinated the investigation, provided resources and mentorship for fieldwork execution and data analysis. PT wrote the original draft. All authors read, edited, gave input, and approved the final manuscript.

## Acknowledgements

We are grateful to all the survey participants. We would like to thank the Department of Tropical Medicine, Tulane University for support. This study was partially supported by the NSF award 1563531; the funding agencies had no involvement in study design, data analysis, or decision to publish.

## Appendix A

Let *x = (E_h_, I_h_, E_g_, I_g_, E_b_, I_b_)^T^* and *dx/dt = F(x) – V(x),* where *F(x)* represents the rate of new infections entering the population, and *V(x) = V^-^(x) – V^+^(x)* represents the rate of movement by other means out of, and into each compartment, respectively.

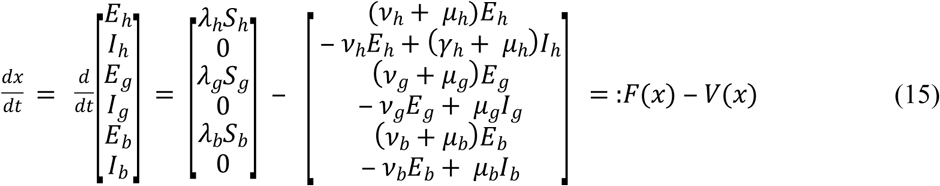

Let *F_0_* and *V_0_* be the Jacobian matrices of the six elements of *F* and *V,* respectively, evaluated at the disease-free equilibrium. Then,

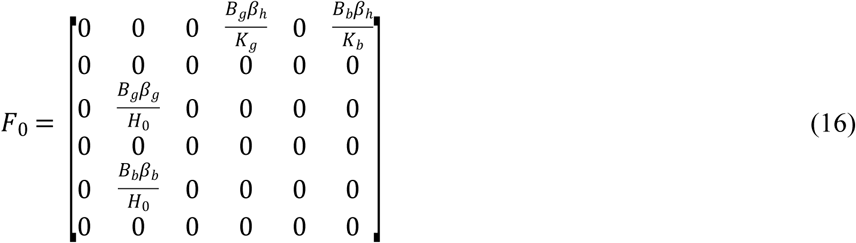

and,

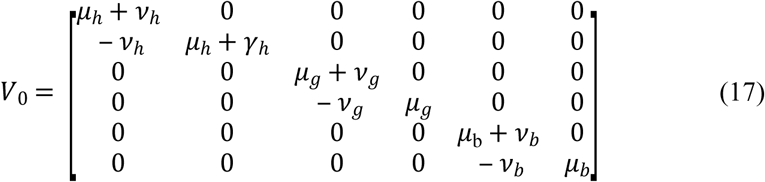

The next generation matrix is

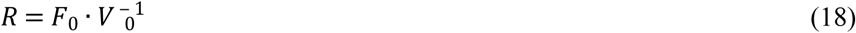

or,

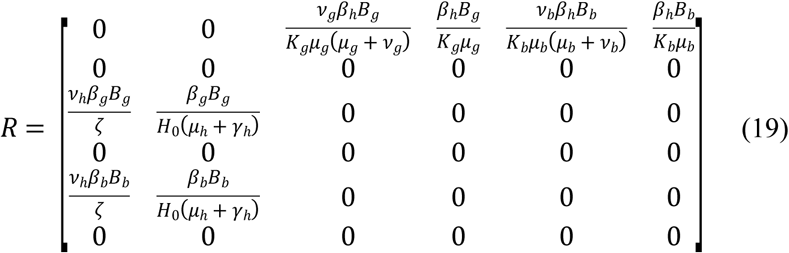

where *ζ* = *H*_0_(*μ_h_* + *ʋ_h_*)(*μ_h_* + *γ_h_*). The *k_ij_*entry of *R* is the average number of cases in class *i* resulting from an infectious individual in class *j*. Finally, R_0_ can be calculated as the absolute value of the largest eigenvalue, or the spectral radius, of the next generation matrix.

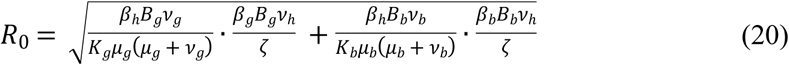

## Supporting Information Legends

**Supplementary Data 1.** The questionnaire used in the preliminary survey.

**Supplementary Data 2.** The questionnaire used in the second survey. The human-mosquito contact rates used in the mathematical model analyses were derived from this questionnaire.

**Supplementary Table 1.** Selected demographic variables of research participants in three study sites.

**Supplementary Figure 1.** A diagram showing the SEIR model compartments and parameters. Black arrows connecting boxes represent the transitions of one disease state to another. Vertical black arrows leaving boxes represent deaths. Vertical black arrows going into boxes represent recruitments of new individuals. Dashed arrows represent contacts between humans and mosquitoes. S_h_, S_g_, and S_b_ represent susceptible human, *Ae. aegypti* and *Ae. albopictus*, respectively. E_h_, E_g_, and E_b_ represent exposed human, *Ae. aegypti*, and *Ae. albopictus,* respectively. I_h_, I_g_, and I_b_, represent infected human, infected *Ae. aegypti*, and *Ae. albopictus*, respectively. R_h_ represent recovered human. Other parameters are described in Table 1 in the main manuscript.

**Supplementary Figure 2.** Bar graphs showing the distributions of research participants’ age and gender in the second survey.

**Supplementary Figure 3.** Graphs showing the average time spent outside in the past 24 hours during weekday and weekend reported by research participants from ORL and TAM. The unit of time duration on the y-axis is in minutes. The time of the day includes ‘morning’ (from 06:00 to 10:00), ‘day’ (from 10:00 to 17:00), ‘evening’ (from 17:00 to 20:00), and ‘night’ (from 20:00 to 06:00 the next day). The error bars represent the standard errors.

**Supplementary Table 2.** Results from a quasi-Poisson regression analysis with a log link function to determine the associations between the response variable, the time spent outside between 5 pm to 6 am reported by research participants, and study sites, weekend/weekday setting, and ages and gender of participants.

**Supplementary Table 3.** Results from a quasi-Poisson regression analysis with a log link function to determine the associations between the response variable, the numbers of mosquito bites reported by research participants, and time spent outside, the month of data collection, study site, and gender of participants.

**Supplementary Figure 4.** The extended sensitivity analysis plots of R_0_ to selected model’s parameters from the DENV transmission model in ORL. Red points represent the baseline values of the parameters.

**Supplementary Figure 5.** The extended sensitivity analysis plots of R_0_ to selected model’s parameters from the DENV transmission model in TAM. Red points represent the baseline values of the parameters.

